# A convergent behavioral-neural profile of parental dysregulation and the link between parent-child brain-state similarity and child negative affect

**DOI:** 10.64898/2026.09.15.751857

**Authors:** Qingyi Li, Ya-Yun Chen, Zexi Zhou, Yang Qu, Jungmeen Kim-Spoon, Tae-Ho Lee

## Abstract

Parent-child neural similarity is often interpreted as a marker of attunement, but its developmental significance may not be uniformly positive across family environments. We tested whether the parent’s own emotional and regulatory profile—measured both behaviorally and neurally—moderates the developmental significance of parent-child neural similarity. Applying a hidden Markov model (HMM-MAR; *K* = 6 states) to whole-brain fMRI timeseries from 32 parent-youth dyads who separately watched the same emotional film, we derived per-dyad measures of brain-state similarity. Parent-child dyads showed greater brain-state similarity than parent-shuffle null pairs (*d* = -0.55, *p* = .004), indicating dyad-specific alignment beyond responses shared across viewers of the same film. Parent self-reported internalizing showed only a weak direct relation to child negative affect (*r* = .31, *p* = .085). However, the association between neural similarity and child negative affect depended on parent internalizing (*b =* 5.62, *p* = .018; *ΔR*² = 15.7%): stronger parent-child similarity predicted higher child negative affect when parents reported higher, but not lower, internalizing. A parallel interaction emerged when parent internalizing was replaced by a neural index of the parent’s own state dynamics (parent neural-state unpredictability, *H*_trans_; *b =* 5.95, *p* = .004), indicating that the effect reflects the parent’s broader regulatory profile rather than a single measurement modality. Both interactions were clearest in the primary model and only partially robust across model orders and parcellations. These findings suggest that dyadic neural similarity should be interpreted in context: resembling a parent with a dysregulated profile may confer risk rather than protection.

## Introduction

A growing body of neuroimaging work has shown that parents and children exhibit measurable similarity in brain activity and connectivity during shared or matched emotional experiences [1–5] (for reviews, see [6, 7]). This similarity has been linked to a range of favorable child outcomes, including emotional competence, daily emotional synchrony, stress regulation, and mental resilience [1, 8–11]. However, emerging evidence suggests that the developmental meaning of neural similarity is not fixed. Zhou et al. [2] examined parent-youth dyads who separately watched the same emotion-evoking animated film and found that neural similarity was associated with more favorable youth’s emotional adjustment only in families with higher cohesion. Similarly, Su et al. [3] linked a more negative family emotional climate to reduced parent-child neural synchrony during naturalistic movie-watching and to higher child internalizing symptoms.

These studies characterize context at the level of the family as a whole, using constructs such as family cohesion and emotional climate. Yet families are composed of individuals, and an overall family-level climate does not imply that each family member functions in the same way [12, 13]. A cohesive family can still include a parent who is highly anxious or emotionally dysregulated, and a child whose brain processes emotional information similarly to that of a calm, well-regulated parent may be in a different developmental situation from one who resembles an anxious or emotionally distressed parent. This points to a more proximal context for interpreting parent-child neural similarity: the parent’s own emotional and regulatory characteristics. This emphasis is consistent with developmental frameworks that conceptualize a parent’s own regulatory characteristics as part of the context in which children’s self-regulation develops [13, 14]. Parents’ internalizing and regulatory characteristics are robustly associated with their children’s emotional development [15–17], yet whether these characteristics also condition what a child’s neural resemblance to the parent means remains unexamined. Together, these perspectives suggest that the developmental significance of parent-child neural similarity may depend on the emotional and regulatory characteristics of the parent whose neural dynamics the child resembles.

Beyond refining the context in which neural similarity is interpreted, a parallel methodological question concerns the temporal resolution at which that similarity is characterized. Prior studies have typically characterized parent-child similarity using regional activation patterns [9] or static functional connectivity pattern [1]. Although informative, these approaches compress complex neural dynamics into a single time-averaged summary. Dynamic functional connectivity methods, such as sliding-window correlation, capture some of this temporal variation, but they rely on a predetermined window length that may not align well with the timescale of the brain’s intrinsic state transitions [18]. An alternative is to model neural dynamics in terms of discrete states whose onset and duration are estimated directly from the data. Hidden Markov models provide such an approach without imposing a fixed temporal window [19, 20], and HMM-derived brain-state synchrony during naturalistic film viewing has recently been shown to differentiate clinical phenotypes of depression [21]. They can therefore identify recurring neural configurations, including transient states that may be obscured by temporal averaging or fixed-window analyses. Accordingly, parent-child neural similarity may extend beyond scan-averaged patterns to the temporal organization of neural responses including how long parents and children occupy particular latent brain states, how often they transition between states, and how predictable those transitions are. Characterizing these dynamics may therefore reveal whether parent-child similarity reflects sustained alignment in neural processing, coordinated shifts between latent states, or both.

Importantly, the HMM further yields an index of how predictable each parent’s own transitions among brain states are during the film (parent neural-state unpredictability, *H*_trans_). Such HMM-derived transition metrics provide reliable indices of individual neural variability and have been linked to behavioral and emotion-regulation measures in developmental samples [19, 22]. This offers a neural characterization of the parent, derived from the same model and grounded directly in brain dynamics rather than self-report. Examining self-report and this neural measure together allows us to test whether the same moderation effect holds across a behavioral and a neural characterization of the parent.

In the present study, we examined whether parents and youth showed dyad-specific similarity in latent brain-state dynamics while separately viewing the same emotion-evoking animated film (Figure 1). Because parents and children were scanned separately, dyadic similarity reflects shared stimulus-evoked processing rather than real-time interpersonal coupling. We analyzed movie-viewing fMRI data from 32 parent-youth dyads, applying a multivariate-autoregressive hidden Markov model (HMM-MAR [19]) to each participant’s brain timeseries. From this model, we derived the two measures described above: dyad-level brain-state similarity, indexing how similarly parents and youth organized their neural responses over time, and each parent’s own neural-state unpredictability (*H*_trans_).

**Figure 1.**
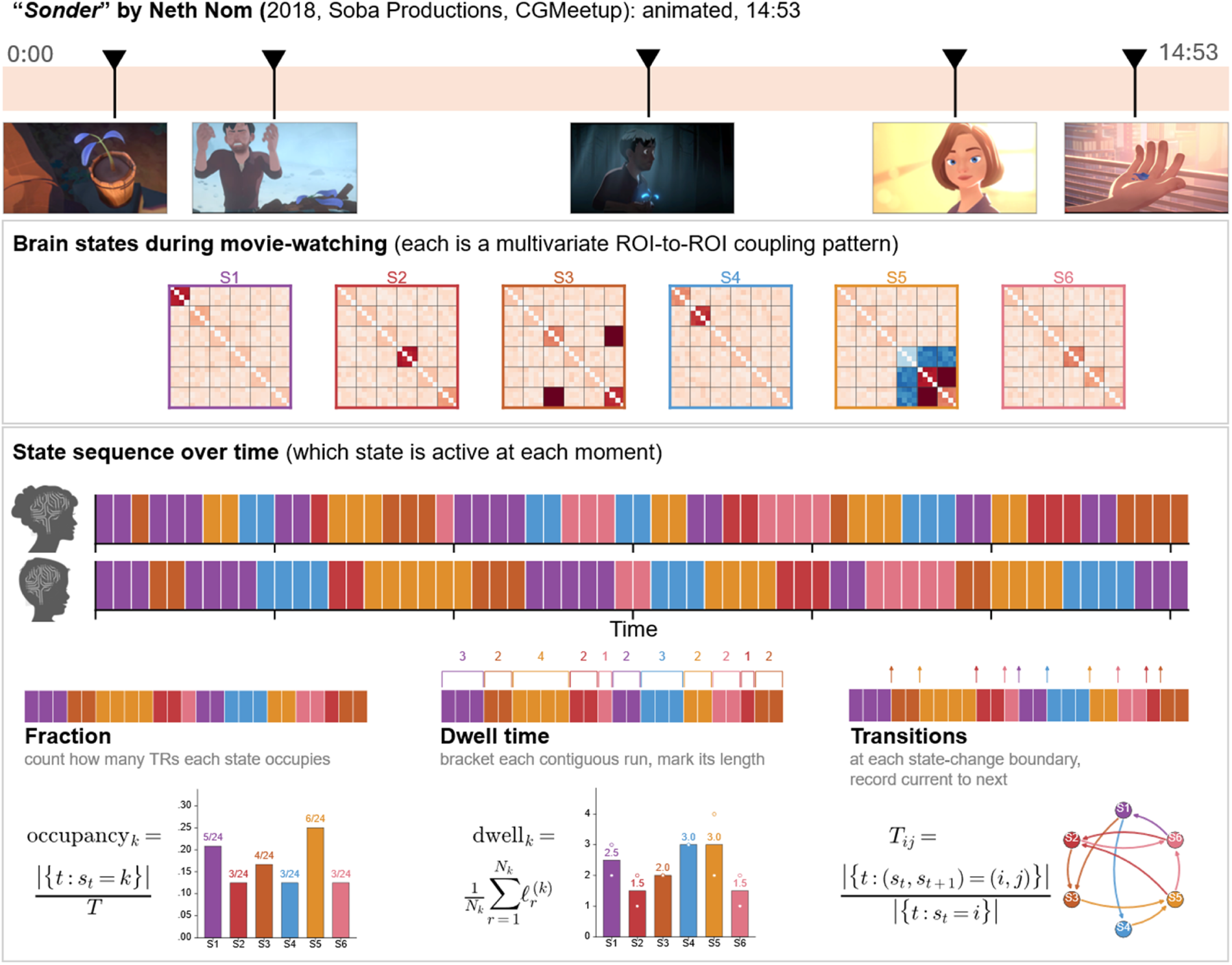
Study design and methods schematic. The animated short *Sonder* [51] (14:53) served as the movie stimulus; parent and child each watched the film at separate scanning sessions, during which approximately 13.1 min (393 volumes) was presented. Each subject’s BOLD time series was fit with HMM-MAR; the Viterbi state path yields per-subject occupancy, dwell-time, and transition metrics, illustrated schematically.

The present study proceeded in three steps. First, we tested whether actual parent-child dyads exhibited greater brain-state similarity than would be expected by chance, establishing whether dyad-specific alignment was present during movie viewing. Second, we tested whether parent internalizing moderated the link between neural similarity and child negative affect—that is, whether the strength or direction of the similarity–negative-affect association depended on the parent’s level of internalizing. Third, we tested whether the same moderation emerged when the parent was characterized neurally by *H*_trans_ rather than by self-report, providing a convergence test based entirely on neural measures. Together, these analyses tested whether parent-child neural similarity carries developmental meaning that depends not only on family-level context, but also on the individual parent’s own emotional and regulatory profile.

## Materials and methods

### Participants and design

The sample comprised 32 parent-youth dyads (32 parents, 32 children): youth ranged in age from 8 to 17 years (M = 11.69, SD = 2.80) and parents ranged from 30 to 64 years (M = 43.53, SD = 7.30). Regarding biological sex, children were 47% female, and parents were 72% female. Regarding race and ethnicity, 69% of youth self-identified as non-Hispanic White American, 16% as Hispanic American, 12% as non-Hispanic Asian American, and 3% as non-Hispanic Black or African American; 81% of parents self-identified as non-Hispanic White American, 13% as non-Hispanic Asian American, 3% as Hispanic American, and 3% as non-Hispanic Black or African American. Thirty of 32 caregivers (94%) were biological parents and 6% were adoptive parents. More details are reported in Table 1. All procedures were approved by the Virginia Tech Institutional Review Board; parents provided informed consent and youth provided assent.

**Table 1.** Sample demographics and descriptive statistics.

|  | Parents (n = 32) | Youth (n = 32) |
| --- | --- | --- |
| **Demographics** |  |  |
| Age, M (SD) | 43.53 (7.30) | 11.69 (2.80) |
| Age range | 30–64 | 8–17 |
| Female, n (%) | 23 (72%) | 15 (47%) |
| **Race/ethnicity** |  |  |
| Non-Hispanic White American | 26 (81%) | 22 (69%) |
| Non-Hispanic Asian American | 4 (13%) | 4 (12%) |
| Hispanic American | 1 (3%) | 5 (16%) |
| Non-Hispanic Black or African American | 1 (3%) | 1 (3%) |
| Biological parent, n (%) | 30 (94%) | — |
| **Self-report measures** |  |  |
| STAI-Trait, M (SD) | 37.00 (9.57) | 34.29 (8.44) |
| STAI-State, M (SD) | 33.28 (9.90) | 32.69 (8.31) |
| CES-D Depression, M (SD) | 9.97 (9.18) | 15.84 (9.46) |
| PANAS-Negative Affect, M (SD) | 24.53 (10.15) | 31.94 (11.53) |
| Internalizing composite, M (SD) | 0.00 (0.89) | — |
| Ego Resilience, M (SD) | 3.77 (0.65) | 3.14 (0.83) |
| **Neural measures (K = 6 movie)** |  |  |
| H<sub>trans</sub>, M (SD) | 0.719 (0.184) | 0.811 (0.212) |

*Note. Self-report measures are reported as sample means (SD); STAI, CES-D, and PANAS are summed totals, whereas Ego Resilience is an item mean. STAI = State-Trait Anxiety Inventory [29]; child STAI-Trait and STAI-State are reported on a harmonized 20–60 metric across age-appropriate forms (STAIC for ages 8–12; adult STAI for ages 13–17). CES-D = Center for Epidemiologic Studies Depression Scale (parent form [30]; child form: CES-DC [52]). PANAS-Negative = 14 negative affect items of the PANAS [31, 32], reported as a summed total (range 14–70). Ego Resilience = 6-item Brief Resilience Scale [36], reported as an item mean (range 1–5), n=27. Internalizing composite = z-mean of parent STAI-Trait, STAI-State, CES-D, and PANAS-Negative Affect ( $\alpha = .92$ ). $H_{trans}$ = Markov entropy rate of HMM state sequence.*

### fMRI data acquisition and analyses

#### Image acquisition

All MRI data were acquired on a Siemens Prisma 3T scanner at Virginia Tech. High-resolution T1 (TR = 2.5 s; TE = 2.06 ms; FA = 8°; 1 mm isotropic voxel; FOV = 256 mm) and T2 (TR = 3.2 s; TE = 563 ms; FA = 120°; 1 mm isotropic voxel; FOV = 256 mm) anatomic images were acquired for tissue segmentation (GM, WM, and CSF mask) and normalization. Functional images for movie watching (393 volumes) were acquired with a gradient-echo echo-planar T2*-weighted imaging sequence (TR = 2 s; TE = 25 ms; FA = 90°; 2.5 x 2.5 mm resolution; 37 interleaved 3.0 mm slices with 0.3 mm gap; FOV = 230 mm; 92 x 92 matrix).

#### Preprocessing

Preprocessing was performed using FSL 6.0.7.10 and ANTs 2.3.5. The first two volumes were discarded for signal equilibration; rigid-body motion was corrected with MCFLIRT; slice timing was corrected with the participant-specific slice-acquisition file; non-brain tissue was removed with BET; data were grand-mean intensity normalized to 10 000; spatial smoothing was applied with a 5 mm FWHM Gaussian kernel; ICA-AROMA was run for denoising of motion-related independent components [23]. ANTs nonlinear registration aligned each participant’s structural to the MNI152 2 mm template and the EPI to the structural via two rigid steps. Excessive head motion was defined as a mean framewise displacement above 0.5 mm; no participant exceeded this criterion, and all were retained.

Three additional steps were applied for the subsequent analysis. First, a 26-column nuisance design was voxel-wise regressed from the preprocessed data within the brain mask. This design included the 24-parameter Friston expansion of the rigid-body motion estimates [24], plus the mean signal from the participant’s CSF and white-matter tissue masks. Second, a fourth-order Butterworth high-pass filter at 0.001 Hz was applied with zero-phase forward-backward filtering. We used this cutoff as a minimal high-pass step intended to reduce ultra-slow drift while avoiding removal of slow fluctuations within the observed scan duration. No low-pass filter was applied, preserving spectral content up to the Nyquist frequency (0.25 Hz at TR = 2 s) for the multivariate-autoregressive model. Third, region-of-interest time series were extracted in each participant’s native EPI space by warping a combined 232-region atlas obtained from the Schaefer 200-cortical parcellation [25] merged with the Tian-2020 Scale-2 32 subcortical atlas [26] with nearest-neighbor interpolation. The unweighted voxel-wise mean within each parcel was taken as that ROI’s time series. The combined atlas was constructed once in MNI 2 mm space, and in 74 voxels of cortex-subcortex adjacency the subcortical (Tian) label was retained. For atlas sensitivity analyses, time series were also extracted using the AAL3 atlas [27] (166 ROIs after excluding ROIs with missing native-space coverage in any participant).

#### HMM-MAR fitting

ROI time series were standardized within each participant (z-scored per ROI) and concatenated across all 64 subjects, yielding a single matrix of size 232 × T_total_, where T_total_ is the total number of concatenated timepoints. A hidden Markov model with multivariate-autoregressive observations (HMM-MAR [19]) was fit in MATLAB using the OHBA-analysis toolbox, with an autoregressive order of 1 and diagonal covariance. The model was fit in PCA space (50 components, 78.2% of variance retained), so individual ROI labels do not enter the fit. For each configuration, three random initializations were run and the fit with the lowest free energy was retained.

We selected K = 6 as the primary model because it returned the lowest free energy (a standard model-fit criterion, where lower values indicate a better fit) across the tested grid of K ∈ {4, 6, 8, 10, 12} (Supplementary Table 1). At K = 6, each of the six states had non-collapsed occupancy with a distinct cortical or cortico-subcortical engagement profile and a distinguishable spectral profile (Figure 2). Per-state functional connectivity was computed as the Pearson correlation matrix across all TRs assigned to each state, pooled over all 64 subjects (232 × 232 for the Schaefer + Tian parcellation). Per-state power spectra (the distribution of signal power across frequencies) were computed across the 0.004 to 0.25 Hz band.

**Figure 2.**
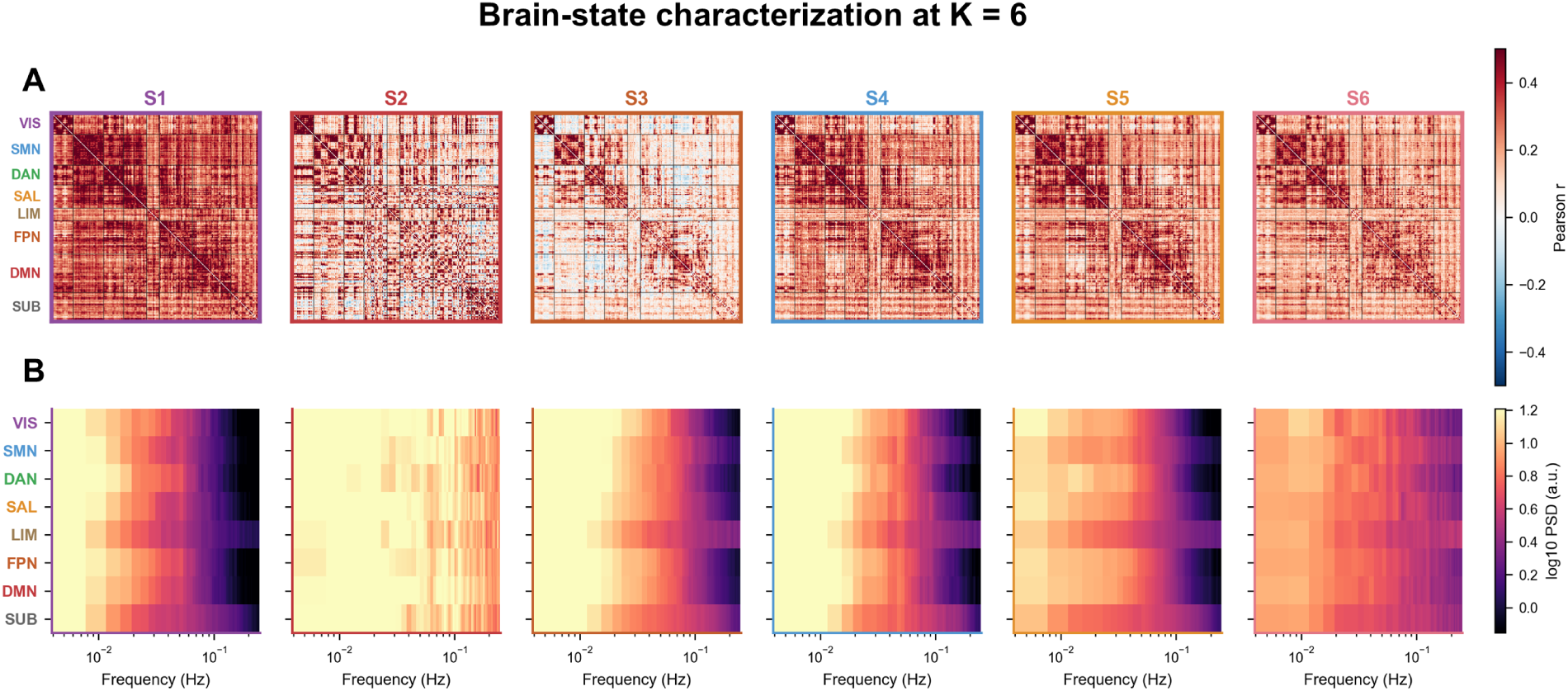
Brain-state characterization at *K* = 6. Each column shows one HMM state (S1 to S6), with the state color used throughout the manuscript on the panel border. (A) Per-state Pearson functional-connectivity matrix across the 232 ROIs (Schaefer-200 cortex + Tian-2020 S2 subcortex), computed on TRs assigned to that state across all 64 subjects. ROI rows and columns are grouped by Yeo-7 cortical networks (VIS, SMN, DAN, SAL, LIM, FPN, DMN) followed by subcortex (SUB) for display only; the HMM is fit on the 232-ROI data after PCA reduction, so network labels do not enter the model. Color scale: Pearson *r* between -0.5 and 0.5. (B) Per-state power spectra, computed as Welch power spectral density on the same state-conditional TRs and averaged within the same eight (Yeo-7 + SUB) groups. Color scale: log₁₀ PSD (a.u.), shared across the six states; x-axis is the 0.004–0.25 Hz band on a log scale.

#### Dyadic brain-state similarity metrics

From each HMM fit, three per-subject indices were extracted: the K-dimensional state-occupancy distribution (fractional time spent in each state), the K × K transition matrix (probability of transitioning from each state to each other state), and the K-dimensional mean dwell-time vector (average run length per state, in seconds). Dyadic similarity was thus quantified in three ways: the symmetric Kullback-Leibler divergence (a measure of how different two probability distributions are; smaller means more similar) between occupancy distributions, the off-diagonal Pearson correlation between transition matrices (larger means more similar; restricting to off-diagonal entries prevents shared state stickiness from inflating dyadic agreement), and the Pearson correlation between dwell-time vectors (larger means more similar).

For group-level inference, a permutation null was generated by randomly re-pairing children and parents from the same cohort (parent-shuffle), with each per-dyad null distribution constructed from 5 000 independent permutations. Occupancy KL divergence was used as the primary similarity index in all subsequent interaction analyses, with its sign reversed so that higher values indicate stronger dyadic alignment.

#### Parent neural-state transition unpredictability

To test whether the interaction pattern also emerged when the parent-level measure was derived from neural dynamics rather than self-report, we computed parent neural-state unpredictability (*H*_trans_) from each parent’s HMM state sequence. Unlike the dyadic similarity measures, which compare parent and child state sequences, *H*_trans_ indexes the predictability of the parent’s own state transitions. To evaluate the construct relevance of H_trans_, we correlated it with 35 additional parent self-report measures available for a subset of 26 parents with complete questionnaire data. The full list of measures and correlations is reported in Supplementary Table 2.

For each parent, the most likely state sequence (Viterbi path) was extracted from the fitted HMM, assigning each time point to its most probable brain state. The K × K transition matrix was estimated from this sequence, per-row Shannon entropies were computed, and these were weighted by the empirical state-occupancy distribution to yield the Markov entropy rate, *H*_trans_ [28], which summarizes in a single number how unpredictable the parent’s moment-to-moment state changes are. Low *H*_trans_ indicates that the next brain state is highly predictable given the current one, while high *H*_trans_ indicates that many alternative transitions are roughly equiprobable.

### Psychological measures

#### Parental internalizing composite profile

Following the intergenerational self-regulation framework [14], we constructed a parent emotional internalizing composite from four parent self-report scales: trait anxiety (STAI-T [29]), state anxiety (STAI-S [29]), depressive symptoms (CES-D [30]), and negative affect (the 14 negative-affect items of the PANAS [31, 32], following Zhou et al. [2]). Each scale score was z-standardized within the cohort, and the four standardized scores were averaged to form the composite (Cronbach’s α = .92). As a specificity check, we constructed a general index of the family relational context from children’s reports on three family subscales—family identity [33], family conflict (reverse-scored [34, 35]), and family obligation [33]. Each subscale was z-scored and averaged, and the composite was entered in place of the parent’s internalizing composite as an alternative moderator.

#### Child outcome variables

The primary child outcome was child recent negative affect measured by the 14-item PANAS [2, 31, 32], indexing recent 1 month emotional experience rather than a stable trait. Two secondary child outcomes were examined to evaluate whether the interaction pattern generalized beyond recent negative affect to related but more stable affective-risk measures: child trait anxiety and child ego resilience [36]. Because participating children spanned 8–17 years and completed age-appropriate anxiety forms with different response scales (3-point STAIC at ages 8–12 [37]; 4-point adult STAI at ages 13–17 [29]), child trait- and state-anxiety scores were linearly rescaled to a common 20–60 metric before analysis. Descriptive statistics are reported in Table 1, and interaction results are in Table 2.

**Table 2.** Interaction model results: parent measure × dyadic neural similarity predicting child PANAS-Negative Affect.

| Level | Outcome | Condition | Parent measure | *b* | *p* | Model *R*<sup>2</sup> | *n* | 95% Bootstrap CI |
| --- | --- | --- | --- | --- | --- | --- | --- | --- |
| Primary | PANAS-Negative Affect | K=6 Schaefer (primary) | Internalizing composite | 5.62 | .018 | .368 | 32 | [0.18, 9.91] |
|  |  | K=6 Schaefer (primary) | *H*<sub>trans</sub> | 5.95 | .004 | .395 | 32 | [1.01, 9.74] |
|  |  | K=4 Schaefer (sensitivity) | Internalizing composite | 4.69 | .076 | .302 | 32 | [-1.18, 9.76] |
|  |  | K=4 Schaefer (sensitivity) | *H*<sub>trans</sub> | 0.92 | .734 | .259 | 32 | [-7.54, 5.68] |
|  |  | K=6 AAL3 (sensitivity) | Internalizing composite | -1.66 | .291 | .259 | 32 | [-6.25, 4.28] |
|  |  | K=6 AAL3 (sensitivity) | *H*<sub>trans</sub> | 4.62 | .038 | .285 | 32 | [-0.25, 10.70] |
|  |  | K=4 AAL3 (sensitivity) | Internalizing composite | 2.58 | .314 | .248 | 32 | [-3.42, 7.70] |
|  |  | K=4 AAL3 (sensitivity) | *H*<sub>trans</sub> | 5.49 | .075 | .258 | 32 | [-0.06, 12.14] |
| Secondary | STAI-Trait anxiety | K=6 Schaefer | Internalizing composite | 3.09 | .072 | .351 | 32 | [-1.32, 5.83] |
|  |  | K=6 Schaefer | *H*<sub>trans</sub> | 1.23 | .445 | .210 | 32 | [-2.58, 4.38] |
|  | Ego Resilience | K=6 Schaefer | Internalizing composite | -0.18 | .456 | .090 | 27 | [-0.60, 0.47] |
|  |  | K=6 Schaefer | *H*<sub>trans</sub> | -0.11 | .567 | .252 | 27 | [-0.50, 0.26] |

*Note.* Grey shading = *p* < .05. K=6 Schaefer composite: $\Delta R^2 = .157$ vs baseline ( $R^2 = .211$ ); LR test $\chi^2(2) = 7.11$ , *p* = .029.

#### Post-scan emotion rating

Immediately after the scan, both parents and children rated their emotional experience during the film on 18 emotion items (e.g., happy, sad, angry, warm, scared, interested) using a 5-point Likert scale (1 = not at all, 5 = extremely). These ratings were used as a behavioral concordance comparator for the neural similarity findings.

### Analytic plan

For each similarity metric, dyad-specificity was tested with a one-sample *t*-test on the per-dyad differences between each dyad’s observed metric and its parent-shuffle null mean. The interaction models were specified as:

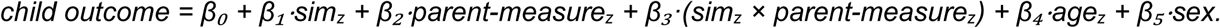

where parent-measure was either the internalizing composite or H_trans_, each tested in a separate model. For each parent measure, a likelihood-ratio test compared a baseline model containing that parent measure and covariates

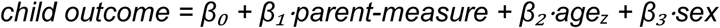

against the corresponding full interaction model, testing whether adding the similarity main effect and its interaction with the parent measure improved model fit. The accompanying change in *R² (ΔR²)* quantified the additional variance explained. To assess the stability of effects under resampling, we computed bootstrap 95% confidence intervals (case resampling, 3 000 resamples) for the key effect sizes and interaction coefficients. Robustness was further evaluated through sensitivity analyses at K = 4 and using the AAL3 atlas (reported in the Results). All models were fit in Python using statsmodels.

## Results

### Latent brain states during movie viewing

The six states (Figure 2) spanned distinct configurations: an occupancy-dominant background state (S5, 42.9% of TRs), two intermediate-occupancy states with cortically organized blocks (S1, 16.9%; S3, 17.2%), and three less frequently occupied states (S2, 1.8%; S4, 10.6%; S6, 10.6%) differing in subcortical engagement and cortico-subcortical coupling. S2 was infrequently occupied, and its per-state characteristics should be interpreted accordingly. Spectral profiles separated two regimes: S1, S3, and S4 showed steep low-frequency-dominant 1/f roll-off, while S5 and S6 displayed flatter spectra with sustained mid-band power.

### Real parent-child pairs show greater dynamic-state similarity in movie viewing

Real parent-child pairs showed smaller state-occupancy KL divergence than parent-shuffle pairs (Figure 3 A1; real mean = 0.768 vs null mean = 1.027; paired *t*(31) = -3.12, *p* = .004; *d* = -0.55, bootstrap 95% CI [-0.99, -0.22]), with 24 of 32 dyads showing lower-than-null KL. Dwell-time vectors were positively correlated on average (Figure 3 A3; mean Pearson *r* = 0.44, bootstrap 95% CI [0.30, 0.58]; median = 0.52; 25 of 32 dyads positive; one-sample *t(*31) = 6.06, *p* < .001, *d* = 1.07, bootstrap 95% CI [0.72, 1.60]). The transition-matrix off-diagonal correlation was in the predicted direction but did not reach significance (Figure 3, A2; real *r* = 0.434 vs null *r* = 0.372; mean difference = 0.06, bootstrap 95% CI [-0.04, 0.16]; paired t(31) = 1.19, p = .25; d = 0.21). For comparison, parent-child concordance in post-scan subjective emotion ratings was only modestly above chance (*Δ* = 0.044), indicating that the dyad-specific alignment captured by the neural measures was not fully reflected in retrospective self-report (Supplementary Figure 1).

**Figure 3.**
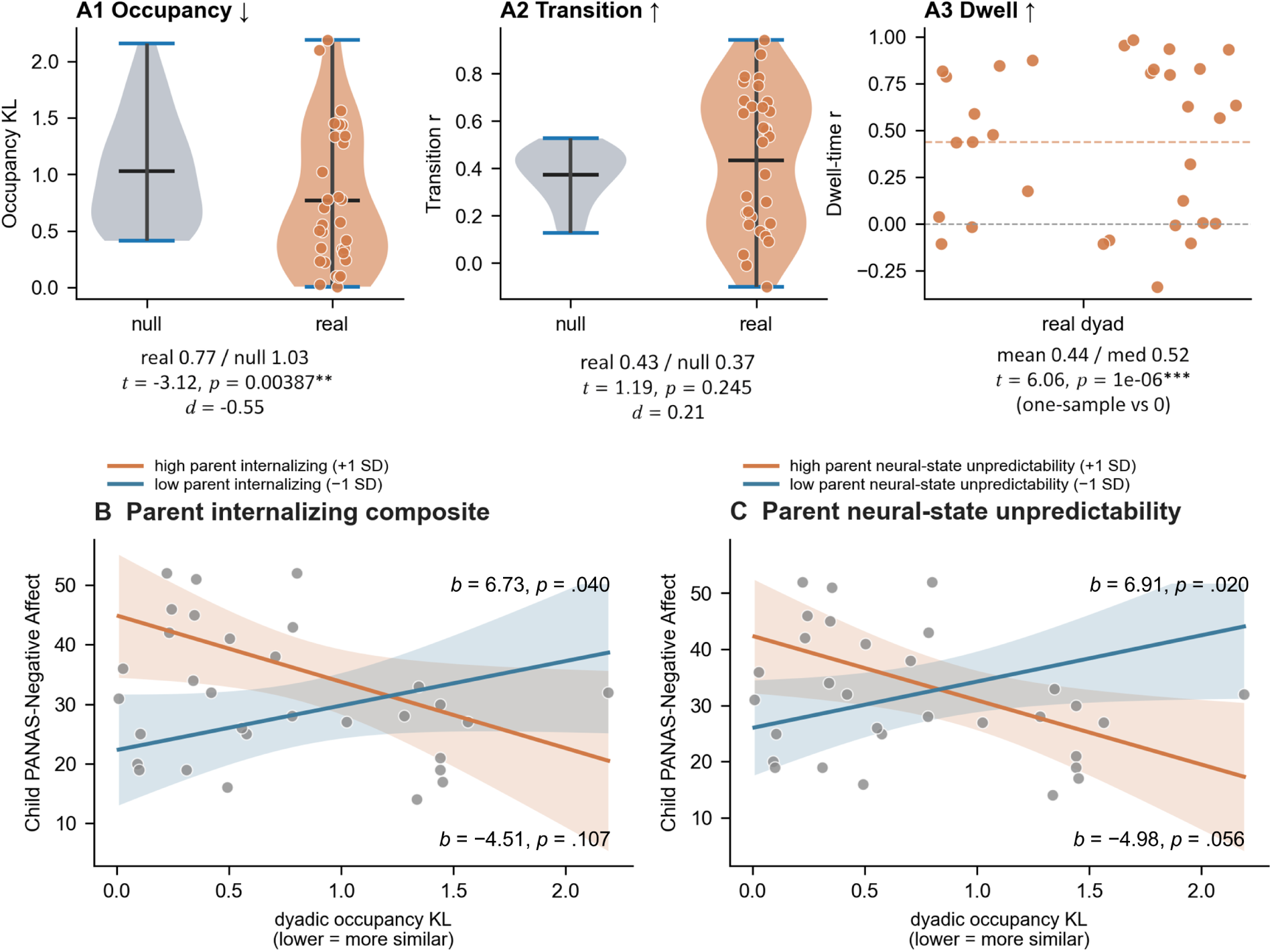
Dyad-specific neural similarity and interaction with parent characteristics. (A) Neural dyadic similarity versus parent-shuffle null (movie K = 6 fit): A1, occupancy KL divergence (d = -0.55, p = .004, bootstrap 95% CI [-0.99, -0.22]); A2, transition-matrix off-diagonal correlation (not significant; mean difference = 0.06, bootstrap 95% CI [-0.04, 0.16]); A3, dwell-time correlation (r = 0.44, p < .001, bootstrap 95% CI [0.30, 0.58]). (B) Parent internalizing composite interaction: predicted child PANAS-Negative Affect as a function of dyadic occupancy KL similarity at high (+1 SD) versus low (−1 SD) parent internalizing, with 95% CI bands (b = 5.62, p = .018, bootstrap 95% CI [0.18, 9.91]). (C) Parent neural-state unpredictability (Htrans) interaction: same format as B, with high (+1 SD) versus low (−1 SD) Htrans (b = 5.95, p = .004, bootstrap 95% CI [1.01, 9.74]).

### Parent internalizing and similarity are not significantly associated with child negative affect

As a baseline, we regressed child recent negative affect on the parent internalizing composite (see Materials and methods), controlling for child age and biological sex. The direct association was not significant (*b =* 2.88, *p* = .156, bootstrap 95% CI [-1.63, 6.50]; zero-order *r* = 0.31, *p* = .085, 95% CI [-0.13, 0.61]). In the full interaction model, the conditional direct association between dyadic similarity and child PANAS-Negative was weak and nonsignificant, b = 1.11, *p* = .564. The corresponding zero-order correlation was likewise weak and nonsignificant, *r* = .11, *p* = .545, bootstrap 95% CI [-.26, .49].

### Parent internalizing moderates the association between parent-child brain-state similarity and child negative affect

We then tested whether parent internalizing and dyadic neural similarity interact in predicting child negative affect. The interaction was significant (*b =* 5.62, *p* = .018, bootstrap 95% CI [0.18, 9.91]; Figure 3B). At high parent internalizing (+1 *SD*), stronger dyadic similarity predicted higher child negative affect, *b* = 6.73, *p* = .040, whereas at low internalizing (−1 *SD*) the association was a nonsignificant negative slope, *b* = -4.51, *p* = .107. The full interaction model explained 36.8% of variance versus 21.1% for the baseline (*ΔR*² = 15.7%; likelihood-ratio test χ²(2) = 7.11, *p* = .029).

### Parent neural-state unpredictability (H*_trans_*) moderates the same association in a similar pattern

To examine whether *H*_trans_ reflects a broader profile of parental dysregulation, we correlated it with 35 parent self-report measures (*n* = 26). These correlations were exploratory and uncorrected, and most were nonsignificant. Among the significant associations, higher *H*_trans_ related to higher trait and state anxiety and nonplanning impulsivity, and to lower fantasy empathy and family identity; the strongest correlation, however, was a positive association with parent-reported influence on the child (r = .53, p = .006), counter to a uniformly maladaptive pattern. Across negative-valence measures, 90% were nominally in the maladaptive direction, but most individual correlations did not reach significance. These exploratory correlations are reported in full in Supplementary Table 2.

Parent *H*_trans_ was not directly associated with child PANAS-Negative in the baseline model, *b* = -1.30, *p* = .524, *R*² = .164; the zero-order correlation was likewise weak and nonsignificant, *r* = -.14, *p* = .446, bootstrap 95% CI [-.49, .28].

We then tested the same interaction model with *H*_trans_ replacing the self-report composite. The interaction was significant for child recent negative affect: *b =* 5.95, *p* = .004, bootstrap 95% CI [1.01, 9.74]. At high parent *H*_trans_ (+1 *SD*), stronger dyadic similarity predicted higher child negative affect, *b* = 6.91, *p* = .020, whereas at low *H*_trans_ (−1 *SD*) the association was a nonsignificant negative slope, *b* = −4.98, *p* = .056 (Figure 3C). The full interaction model explained 39.5% of variance versus 16.4% for the baseline (*ΔR*² = 23.1%; likelihood-ratio test χ²(2) = 10.34, *p* = .006).

### Robustness checks

#### Outcome specificity

When the same interaction models were applied to alternative child outcomes, the interactions were in the same direction but did not reach significance. For the parent internalizing composite: child trait anxiety: *b =* 3.09, *p* = .072, bootstrap 95% CI [-1.32, 5.83]; child ego resilience: *b =* -0.18, *p* = .456, bootstrap 95% CI [-0.60, 0.47]. For *H*_trans_: child trait anxiety: *b =* 1.23, *p* = .445, bootstrap 95% CI [-2.58, 4.38]; child ego resilience: *b =* -0.11, *p* = .567, bootstrap 95% CI [-0.50, 0.26]. The interaction was therefore specific to child recent negative affect in the present sample. Moreover, the general index of the family relational context did not moderate the similarity effect (*b* = 0.46, 95% CI [-3.48, 4.40], *p* = .81), despite a substantial association with child negative affect (*r* = -.49) and a near-zero correlation with neural similarity (*r* = -.04). The moderation was thus specific to the parent’s own internalizing, not the broader family relational context.

#### Sensitivity to K

To test whether the interaction pattern depends on the choice of *K* = 6, we re-ran the full pipeline at *K* = 4 using the same variables as the primary analysis. The dyad-versus-null occupancy KL contrast remained significant (paired *t*(31) = -2.10, *p* = .044, *d* = -0.37, bootstrap 95% CI [-0.77, -0.04]). The composite interaction was in the same direction as the K = 6 result (*b =* 4.69, *p* = .076, bootstrap 95% CI [-1.18, 9.76]) but did not reach significance. The *H*_trans_ interaction was not significant at *K* = 4 (*b* = 0.92, *p = .734, bootstrap* 95% CI [-7.54, 5.68]). The primary interaction pattern is therefore directionally consistent at *K* = 4 but weaker.

#### Sensitivity to atlas

To ensure findings were not specific to the Schaefer-200 + Tian-S2 parcellation, we re-extracted time series with the AAL3 atlas [27] and re-fit the HMM-MAR at *K* = 4 and *K* = 6 using the same variables as the primary analysis. Dyad-versus-null occupancy KL was significant for both AAL3 *K* = 6 (*t*(31) = -4.24, *p* < .001, *d* = -0.75, bootstrap 95% CI [-1.04, -0.52]) and AAL3 *K* = 4 (*t*(31) = -3.70, *p* < .001, *d* = -0.65, bootstrap 95% CI [-0.89, -0.46]), indicating that dyad-specific alignment is robust across parcellations. The *H*_trans_ interaction was significant on the AAL3 atlas at *K* = 6, although its bootstrap CI included zero (*b =* 4.62, *p* = .038, bootstrap 95% CI [-0.25, 10.70]) and was marginal at *K* = 4 (*b =* 5.49, *p* = .075, bootstrap 95% CI [-0.06, 12.14]). The composite interaction, however, did not replicate on the AAL3 atlas (*K* = 6, *b =* -1.66, *p* = .291, bootstrap 95% CI [-6.25, 4.28]; *K* = 4, *b =* 2.58, *p* = .314, bootstrap 95% CI [-3.42, 7.70]). The greater cross-atlas consistency of *H*_trans_ relative to the composite is consistent with the fact that *H*_trans_ and dyadic similarity are both derived from the same HMM decomposition, whereas the composite is an external questionnaire measure whose interaction with similarity may be more sensitive to how similarity is operationalized across parcellations. Nevertheless, parent H*_trans_* and dyadic similarity were only weakly and nonsignificantly correlated (r = -.13, p = .50), indicating that the two HMM-derived measures are not redundant.

## Discussion

The present study examined whether the association between parent-child brain-state similarity and child negative affect depends on the parent’s own regulatory profile, characterized both behaviorally and neurally. Three findings were observed. First, real parent-child dyads showed greater similarity in latent brain-state organization than random pairs, particularly in how much time they spent in each brain state; parent and child state dwell times were also positively correlated. Second, parent internalizing alone was not significantly associated with child recent negative affect, but parent internalizing significantly interacted with dyadic brain-state similarity: stronger dyadic similarity was associated with higher child negative affect when parents reported higher internalizing, but not when parents reported lower internalizing. Third, the same pattern was found when parent self-reported internalizing was replaced by a purely neural measure of how unpredictable the parent’s own brain-state transitions were during the film, derived from the same model as the dyadic similarity measure.

This pattern is consistent with the framework proposed by Bridgett et al. [14], in which the parent’s own regulatory characteristics serve as a developmental context for the child’s emerging self-regulation. The interaction model explained 15.7% more variance in child recent negative affect than the baseline model (parent self-report, age, and sex), suggesting that brain-state similarity provides additional information about which dyads show the strongest association between parent internalizing and child recent negative affect. The specificity to recent negative affect, with only directionally consistent effects for trait anxiety and ego resilience, may reflect temporal alignment between measures, as recent negative affect captures relatively proximal emotional experience, whereas trait anxiety and ego resilience reflect more stable, trait-like characteristics that may be less sensitive to state-level parent-child neural similarity. One possibility is that the children in this cohort (ages 8–17) are within a period in which prefrontal regulatory systems are still maturing [38–41]. Similarity to a parent whose affective processing is characterized by elevated anxiety and negative affect may relate to a form of shared processing that is less well suited to the child’s developing regulatory capacity. This interpretation is compatible with the observed pattern but remains one interpretation among several. The framework thus provides an interpretive context rather than evidence of transmission; the design cannot rule out shared genetic, temperamental, or environmental explanations.

These findings add to the understanding of parent-child neural similarity in two complementary ways—in what is measured and in how its meaning is conditioned. Prior work has begun to qualify the link between parent-child neural similarity and favorable child outcomes [1, 8–11] by relational context. In this same sample, Zhou et al. [2] found that the association between connectivity-pattern similarity and youth emotional adjustment was strongest in higher-cohesion families. Notably, the general index of the family relational context did not moderate the present HMM-based similarity effects; because this index differed from the cohesion measure used previously, whether contextual moderation depends on the level at which neural similarity is characterized remains to be tested directly. We extend this work by identifying the parent’s own affective characteristics as another conditioning context: greater parent-child neural similarity predicted higher child recent negative affect when parent internalizing was high, whereas no significant association was observed when parent internalizing was low. Thus, beyond whether a family is cohesive, it may matter whom specifically the child resembles. This finer moderation was visible because we characterized similarity in latent brain-state dynamics rather than scan-averaged connectivity: how a parent organizes and moves through brain states over time carries conditional information that time-averaged measures may obscure.

The interaction between dyadic similarity and parent internalizing held when parent self-reported internalizing was replaced by *H*_trans_. This reduces the likelihood that the interaction depends on crossing questionnaires with brain data, because *H*_trans_ is derived entirely from the parent’s neural data. Exploratory correlations (Supplementary Table 2) raised the possibility that *H*_trans_ relates to emotional and regulatory characteristics beyond internalizing, but they were uncorrected, mixed in direction, and based on 26 parents. The convergence between questionnaire-based internalizing and *H*_trans_ nevertheless suggests that the interaction may reflect meaningful parent characteristics rather than a feature unique to one measurement modality.

Several features of the design constrain interpretation. First, the sample included only 32 dyads, limiting the precision of interaction estimates and preventing reliable identification of which dimensions of parent internalizing drive the effect. Second, the cross-sectional design cannot determine whether parent-child neural similarity is an antecedent, consequence, or co-determinant of child adjustment, nor can it distinguish among possible sources such as socialization, shared environment, or common temperament. Third, the findings were based on a single emotional film and may not generalize to other stimuli or contexts. Fourth, dynamic functional connectivity remains a methodologically active area [18, 42–46]. The HMM-MAR pipeline addressed several concerns through generative state-space estimation and explicit model selection [47, 48], with *K* = 6 selected as the lowest-free-energy solution across K = 4, 6, 8, 10, and 12, although its free-energy advantage over K = 4 was small (approximately 0.04%). Nevertheless, support across sensitivity analyses was partial rather than uniform. Neither interaction was significant across all model orders and parcellations, and the pattern was clearest in the primary K = 6 Schaefer + Tian model. Finally, child anxiety was assessed with age-appropriate forms (3-point STAIC at ages 8–12; 4-point adult STAI at ages 13–17) linearly rescaled to a common metric; because the form boundary coincides with age, this harmonization is imperfectly separable from child age.

Several directions follow from these findings. The specificity to recent negative affect raises the possibility that the developmental meaning of parent-child neural similarity is especially relevant to ongoing emotional fluctuations. Ecological momentary assessment could test whether similarity in parents’ and children’s daily affective patterns also varies according to the parent’s emotional and regulatory characteristics, thereby providing complementary evidence at a different timescale. Larger cohorts, including clinical and high-risk samples, would permit more reliable estimation of interactions and clearer decomposition of the parental dimensions involved. Extending parent-level measures beyond internalizing symptoms to include parenting behavior and family predictability [49, 50] may further clarify which parental characteristics are most consequential. If conditional patterns are also observed in daily life, ecological momentary interventions targeting parental emotional states during periods of heightened distress could eventually be explored. Overall, the findings suggest that parent-child similarity in latent brain-state dynamics is not uniformly associated with child outcomes; its developmental significance may depend on the emotional and regulatory characteristics that the parent brings to the dyad.

## Acknowledgements

We thank the former and current members of the Affective NeuroDynamics & Development Lab at Virginia Tech for their help with data collection. We are grateful to the adolescents and parents who participated in our study. This work was supported by a Virginia Tech Institute for Society, Culture and Environment research award to T.-H.L.

## Author contributions

T.-H.L. designed research; Y.-Y.C. performed research; Y.-Y.C., Z.Z. and Y.Q. contributed new reagents or analytic tools; Q.L. and T.-H.L. analyzed data, interpreted the results and wrote the paper; and Y.-Y.C., J.K.-S., Z.Z. and Y.Q. edited the manuscript. All authors approved the final version.

## Data availability

Data are available on the Open Science Framework at https://osf.io/j6qvc. Raw structural MRI images are not publicly available because the consent obtained from participating families does not permit public release of individual brain images.

## Code availability

The analysis code is available on the Open Science Framework at https://osf.io/j6qvc.

## Use of generative AI

Claude (Opus 5; Anthropic) was used during manuscript preparation for language editing and consistency checking, and to draft portions of the analysis code, which the authors reviewed and verified. It was not used to design the study, to collect or generate data, or to interpret results. The authors take full responsibility for the content.

