## Supplementary Materials for "A convergent behavioral-neural profile of parental dysregulation and the link between parent-child brain-state similarity and child negative affect"

### Supplementary Table 1. Model selection: free energy across K values

| K | AR order (p) | Free energy | Effective K |
|---|---|---|---|
| 4 | 1 | 1 798 839 | 4 |
| **6** | **1** | **1 798 068** | **6** |
| 8 | 1 | 1 799 737 | 8 |
| 10 | 1 | 1 802 782 | 10 |
| 12 | 1 | 1 808 529 | 12 |

*Note. HMM-MAR fit to concatenated movie-watching fMRI timeseries of 64 subjects (32 parents, 32 children) with diagonal covariance, AR order p = 1, PCA reduction to 50 components. Three random initializations per K; lowest free energy retained. Lower free energy = better fit. Grey/bold = selected model (K = 6).*

**Supplementary Table 2. Exploratory correlations between parent neural-state unpredictability ($H_{trans}$) and parent self-report measures**

| Measure | Valence | *r* | *p* | n | 95% Bootstrap CI |
|---|---|---|---|---|---|
| **Negative-valence measures** | | | | | |
| Trait anxiety | negative | +0.440 | 0.025 | 26 | [-0.02, 0.78] |
| State anxiety | negative | +0.404 | 0.041 | 26 | [-0.08, 0.73] |
| Depression | negative | +0.169 | 0.408 | 26 | [-0.36, 0.56] |
| PANAS-negative | negative | +0.236 | 0.246 | 26 | [-0.24, 0.60] |
| Impulsivity (total) | negative | +0.277 | 0.170 | 26 | [-0.16, 0.63] |
| Attentional impulsivity | negative | -0.008 | 0.969 | 26 | [-0.37, 0.36] |
| Motor impulsivity | negative | +0.204 | 0.317 | 26 | [-0.26, 0.57] |
| Nonplanning impulsivity | negative | +0.390 | 0.049 | 26 | [-0.07, 0.70] |
| BIS (behavioral inhibition) | negative | +0.013 | 0.951 | 26 | [-0.30, 0.40] |
| Sensation seeking | negative | +0.286 | 0.157 | 26 | [-0.16, 0.57] |
| DERS total | negative | +0.152 | 0.459 | 26 | [-0.27, 0.53] |
| DERS Awareness | negative | +0.208 | 0.308 | 26 | [-0.25, 0.55] |
| DERS Clarity | negative | +0.252 | 0.215 | 26 | [-0.37, 0.66] |
| DERS Goals | negative | -0.198 | 0.333 | 26 | [-0.57, 0.25] |
| DERS Impulse | negative | +0.021 | 0.919 | 26 | [-0.41, 0.37] |
| DERS Nonaccept | negative | +0.116 | 0.572 | 26 | [-0.20, 0.42] |
| DERS Strategies | negative | +0.249 | 0.220 | 26 | [-0.17, 0.54] |
| ER Suppression | negative | +0.335 | 0.101 | 25 | [-0.16, 0.70] |
| Controlling parenting | negative | +0.117 | 0.568 | 26 | [-0.39, 0.47] |
| Perceived stress | negative | +0.088 | 0.668 | 26 | [-0.46, 0.62] |
| Personal distress | negative | +0.246 | 0.237 | 25 | [-0.11, 0.55] |
| **Positive-valence measures** | | | | | |
| PANAS-positive | positive | +0.058 | 0.777 | 26 | [-0.42, 0.58] |
| Self-control | positive | -0.037 | 0.858 | 26 | [-0.53, 0.39] |

| | | | | | |
|---|---|---|---|---|---|
| Cognitive reappraisal | positive | +0.059 | 0.776 | 26 | [-0.42, 0.55] |
| Autonomy support | positive | -0.386 | 0.051 | 26 | [-0.72, 0.03] |
| PSE: Parent influence | positive | +0.525 | 0.006 | 26 | [0.29, 0.72] |
| PSE: Developmental | positive | +0.008 | 0.968 | 26 | [-0.41, 0.45] |
| Empathy total (IRI) | positive | -0.275 | 0.184 | 25 | [-0.66, 0.19] |
| Perspective taking | positive | -0.221 | 0.287 | 25 | [-0.66, 0.29] |
| Empathic concern | positive | -0.258 | 0.214 | 25 | [-0.56, 0.14] |
| Fantasy empathy | positive | -0.459 | 0.021 | 25 | [-0.73, -0.07] |
| Family Relationship | positive | -0.213 | 0.306 | 25 | [-0.59, 0.27] |
| Family Identity | positive | -0.424 | 0.035 | 25 | [-0.69, -0.13] |
| BAS approach | positive | +0.188 | 0.358 | 26 | [-0.26, 0.55] |
| Prosocial behavior | positive | -0.274 | 0.186 | 25 | [-0.58, 0.20] |

*Note.* $H_{trans}$ *= Markov entropy rate (K=6, movie, Schaefer). Grey = p < .05 uncorrected. Neg-valence: 19/21 (90%) in maladaptive direction. Pos-valence: 9/14 (64%) in adaptive direction. All correlations exploratory, uncorrected. Impulsivity subscales were assessed with the Barratt Impulsiveness Scale [1]; empathy subscales with the Interpersonal Reactivity Index [2]*; and emotion suppression and cognitive reappraisal with the Emotion Regulation Questionnaire [3].

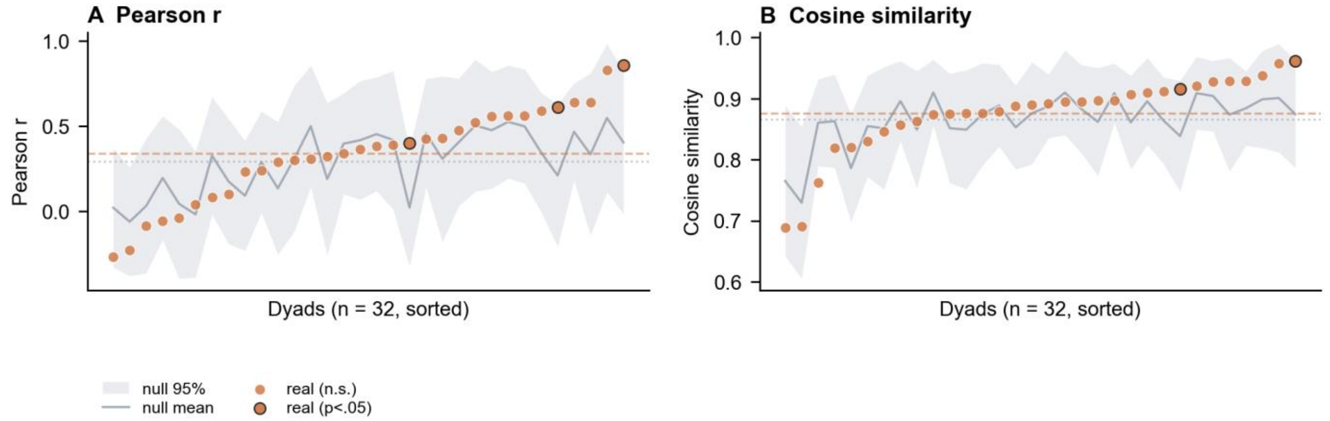


**Supplementary Figure 1.** Behavioral parent-child concordance from post-scan emotion ratings. Per-dyad similarity (Pearson r and cosine similarity) of the 18-item emotion rating vector between parent and child, plotted against the parent-shuffle null distribution. Dyads are sorted by real similarity value. Real parent-child concordance was reliably but modestly above chance (group-mean Pearson r = 0.339, delta versus null = 0.044; 27 of 32 dyads positive). This contrast suggests that the neural approach captures aspects of shared parent-child processing that are not fully reflected in retrospective subjective ratings, which summarize the overall emotional experience rather than the temporal organization of neural responses during the film.
